# Localization and function of key axonemal microtubule inner proteins and dynein docking complex members reveal extensive diversity among vertebrate motile cilia

**DOI:** 10.1101/2024.02.14.580281

**Authors:** Hao Lu, Wang Kyaw Twan, Yayoi Ikawa, Vani Prem Khare, Ishita Mukherjee, Kai Xin Chua, Saikat Chakrabarti, Hiroshi Hamada, Sudipto Roy

## Abstract

Vertebrate motile cilia are broadly classified as (9+2) or (9+0), based on the presence or absence of the central pair apparatus, respectively. Cryogenic electron microscopy (cryo-EM) analyses of (9+2) cilia have uncovered an elaborate axonemal protein composition; whether these features are relevant to (9+0) cilia remain unclear. We previously demonstrated that Cfap53, a key microtubule inner protein (MIP) as well as centriolar-satellites component, is essential for motility of (9+0), but not (9+2) cilia. Here, we show that in (9+2) cilia, Cfap53 functions redundantly with a paralogous MIP, Mns1. Mns1 localizes to ciliary axonemes, and combined loss of both proteins in zebrafish and mice, caused severe loss of outer dynein arms (ODAs) of (9+2) cilia, significantly affecting their motility. Moreover, using immunoprecipitation, we demonstrate that while Mns1 can self-associate and interact with Cfap53, Cfap53 is unable to self-associate. Finally, we show that multiple additional dynein interacting proteins, such as the ODA docking complex (ODA-DC) members, show strikingly distinct localization patterns between various motile cilia-types. Our findings clarify how paralogous MIPs, Cfap53 and Mns1, function in regulating motility of (9+2) versus (9+0) cilia, and establish that localization pattern of other key motility proteins also differ between these cilia-types, further emphasizing extensive structural variations among these organelles.

## Introduction

Motile cilia are biological nanomachines, capable of rhythmic beating for circulation of fluids over epithelia or locomotion of individual cells and organisms. For example, motile cilia lining the airways and the brain ventricles of mammals function in clearing mucus and circulation of cerebrospinal fluid (CSF), respectively, while motile cilia on protozoans, larval forms of invertebrates and sperm are required for propulsion through fluid medium ^1^. In humans, defective motile cilia result in a number of pathological consequences ranging from respiratory disease, hydrocephalus (expansion of the brain ventricular cavities due to impaired CSF flow) and infertility, collectively called primary ciliary dyskinesia (PCD) ^2^. The core or axoneme of a prototypical motile cilium consists of a circular arrangement of 9 peripheral microtubule doublets (a complete A tubule associated with an incomplete B tubule) that surround a pair of singlet microtubules located at the centre along with their associated structures, like the radial spokes - the classic (9+2) arrangement (**Figure 1A**). Inner dynein arms (IDAs) and ODAs are anchored to the peripheral microtubules and are responsible for the beating activity of the organelle. One important variation on this (9+2) configuration are cilia with a (9+0) arrangement of microtubules, lacking the central pair apparatus and its associated structures, like the radial spokes (**Figure 1B**). Loss of the central pair and the radial spokes is believed to allow rotational movement of the (9+0) cilia, in contrast to the whiplash-like, back-and-forth planar beating of the (9+2) cilia. This idea has emerged primarily from the observation that loss of the central pair from otherwise (9+2) cilia confers on them a rotary beat pattern ^3^. The best characterized (9+0) cilia are found within the left-right organizer (LRO) in embryos of several vertebrate species (such as Kupffer’s vesicle (KV) of teleost fishes and the ventral node of the mouse embryo), where their rotary motion is essential for the vectorial flow of extraembryonic fluid within the LRO cavity that initiates asymmetric development of visceral organs ^1^. Consequently, PCD patients are also often afflicted with mirror image reversal (*situs inversus*) or randomization (*situs ambiguus* or heterotaxy) in the disposition of visceral organs, presumably due to dysfunction of their LRO cilia ^2^.

**Figure 1.**
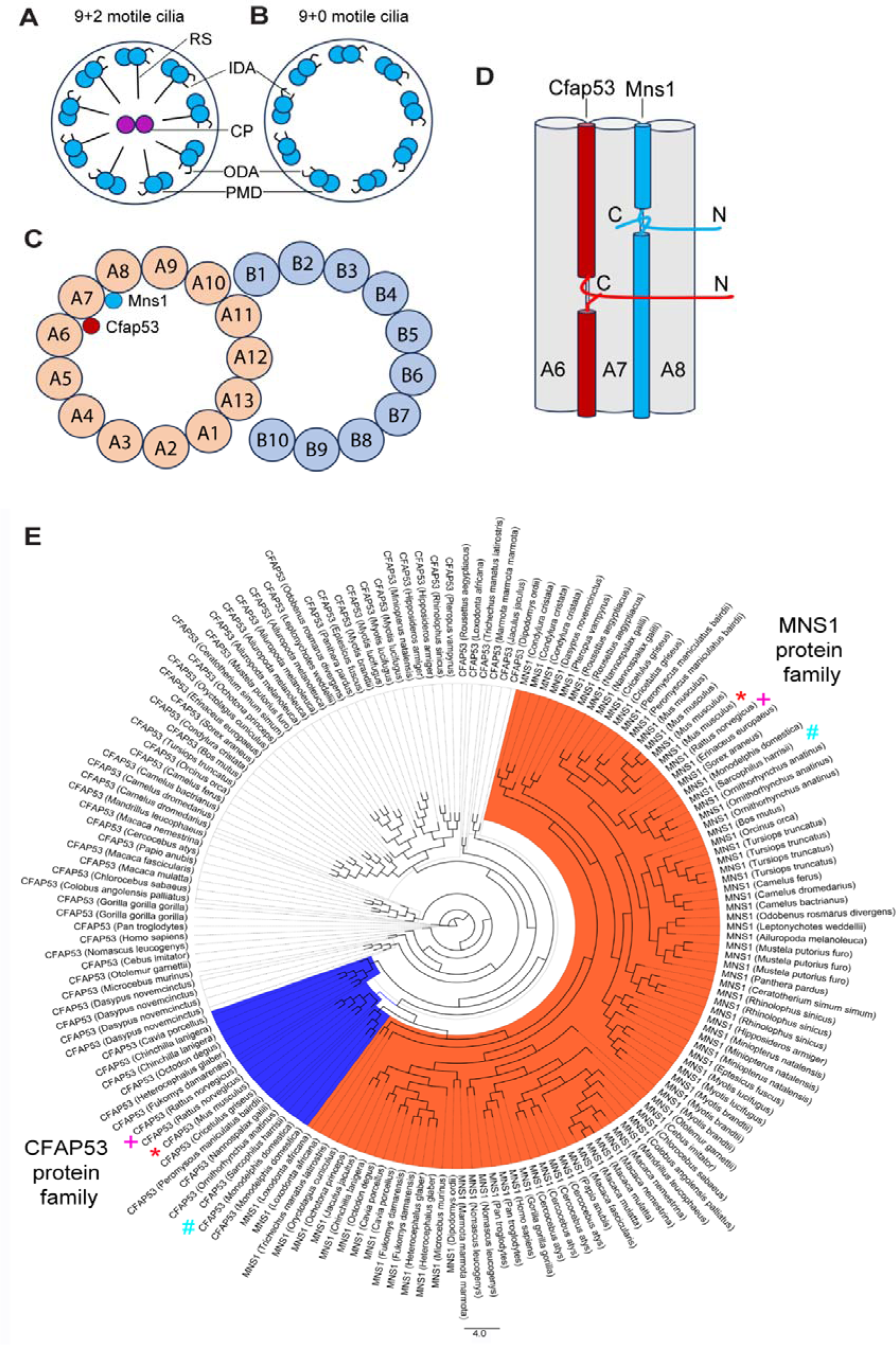
Schematics of types of motile cilia, location of Cfap53 and Mns1 in the ciliary axoneme and phylogenetic relationship of Cfap53 and Mns1 proteins. (A) Schematic diagram of the structure of the (9+2) motile cilium. (B) Schematic diagram of the structure of the (9+0) motile cilium. IDA = inner dynein arm, ODA = outer dynein arm, RS = radial spokes, CP = central pair, PMD = peripheral microtubule doublets. (C) Schematic transverse section through a peripheral microtubule doublet showing Mns1 and Cfap53 localization within A tubule. (D) Schematic longitudinal section through the A tubule showing location of Cfap53 and Mns1 long the grooves of A6 and A7 and A7 and A8 protofilaments, respectively. N = N-terminus, C = C-terminus. (E) Phylogenetic relationship of Mns1 and Cfap53 close orthologs. Cfap53 and Mns1 proteins in *Rattus* sp. [+], *Mus* sp. [*] and some other species [#] cluster together as shown in the phylogenetic tree.

Cryo-EM has revolutionized our understanding of the structural organization of cilia. Studies of motile ciliary axonemes from a wide variety of sources, such as the alga *Chlamydomonas*, bovine and human airways as well as sea urchin and mammalian sperm have not only provided us with a reasonably detailed molecular map of the different proteins and protein complexes that constitute the motility apparatus (such as the dynein arms, radial spokes and the nexin-dynein regulatory complex), but also those that decorate the outer and inner surfaces of the axonemal microtubules ^4–9^. These microtubule-associated proteins are thought to facilitate the anchoring of important motility proteins such as the dynein arm complexes to the microtubules as well as provide structural strength to the microtubules so that they are able to withstand the deformation forces generated during motility. While these data underscore a largely conserved axonemal architecture from protozoans to mammals, they have also revealed notable differences between species (for example, between *Chlamydomonas* and mammals) and well as cilia from different regions of the same organism (for example, respiratory cilia versus sperm flagella from mammals) ^4–8^. However, it is important to note that all of this information pertains exclusively to (9+2) cilia as they are amenable to large-scale isolation in a relatively pure form, such as through deciliation of algal cells and sperm or the trachea from slaughtered mammals. By contrast, the (9+0) motile cilia are relatively limited in numbers and are located in tissues and organs that make their isolation difficult (in the ventral node of the mouse embryo, where these cilia have been best characterised, they number between 200-300 ^10^), thereby precluding ultrastructural characterisation using current technology. Thus, this leaves open the question whether the structural organization of ciliary proteins vary between motile cilia with (9+0) versus (9+2) axonemes, and if so, to what degree.

Functional studies with ciliary proteins are indeed indicating that the localization pattern and motility requirements for even highly evolutionarily conserved components can be rather different between the two kinds of cilia. For instance, Cafp53, is a helical protein and a major filamentous MIP within the A tubule (in the groove between protofilaments A6 and A7) of cilia from protozoans to mammals, as revealed by cryo-EM (**Figure 1C,D**) ^4,5^. Consistent with this, immunolocalization with antibodies to the endogenous protein as well as epitope-tagged functional variants have shown that the protein localizes along the axonemes of (9+2) cilia in the zebrafish, mice and humans ^11,12^. Intriguingly, Cfap53 is also a centriolar-satellites protein in zebrafish and mice associated with the basal bodies of (9+2) cilia, and in (9+0) cilia, it was found to localize exclusively to this compartment (zebrafish KV cilia) or preferentially to this compartment and much less along the axonemes (mouse nodal cilia) ^11–13^. Furthermore, loss-of-function studies with the zebrafish and mice have shown that while the (9+0) LRO cilia almost completely lost their motility, (9+2) cilia remained motile, albeit with some alterations to their beat frequency and amplitude ^11,12,14^. These defects in ciliary motility ensue from a near complete (from (9+0) cilia) or partial loss (from (9+2) cilia) of the ODAs, consistent with biochemical and cryo-EM data implicating association of Cfap53 (either directly or through linker MIPs) with the ODA-DC member Odad4, as well as several of the ODA dynein proteins themselves ^5,11,12^. In line with these molecular data, patients with mutations in the *CFAP53* gene exhibit strong laterality defects signifying disruption of LRO cilia motility, but have a very mild alteration to the motility pattern of their respiratory (9+2) cilia ^11,14,15^.

We were intrigued by the differential localization and function of Cfap53 in the two cilia-types and set out to investigate why the protein is largely dispensable for motility of the (9+2) cilia. Here, we present evidence that Cfap53 is paralogous to another evolutionarily conserved filamentous MIP present within the A tubule, Mns1, that binds the groove between protofilaments A7 and A8, just adjacent to A6 and A7 occupied by Cfap53 (**Figure 1C,D**) ^4,5^. We found that unlike Cfap53, which is also a centriolar-satellites protein, Mns1 localizes mostly to the axonemes of both cilia-types in the zebrafish as well as mice. However, like Cfap53, loss of Mns1 from both organisms significantly impacted the motility of cilia within the LRO, but the motility of (9+2) cilia was much less severely affected. Consistent with an overlapping function of the two paralogues in (9+2) cilia, in animals deficient in both proteins, these cilia almost completely lost all ODAs in mice and showed a range of motility defects, from highly abnormal beating to total paralysis, in the zebrafish. Finally, to further explore the molecular differences between (9+2) and (9+0) cilia, we also examined the localization of 3 highly conserved and core ODA-DC members - Odad1, -3 and -4, to (9+0) and (9+2) cilia of the zebrafish embryo. Surprisingly, we found that while all the Odad proteins exhibited axonemal localization in (9+2) cilia, Odad1 and 3 localized exclusively around the basal bodies of (9+0) cilia, while Odad4 was present along their axonemes, respectively.

Thus, using genetic and cell biological analysis with the zebrafish and mice, we have clarified how two paralogous MIPs, Cfap53 and Mns1, function in regulating the motility of (9+2) and (9+0) cilia. Since mutations in the human orthologs for both genes have been implicated in ciliopathies with the manifestation of male infertility and heterotaxy ^11,14–17^, our data provide valuable mechanistic framework for understanding the pathobiology of these disease phenotypes. Furthermore, our protein localization studies with core ODA-DC members offer additional evidence for extensive molecular distinctiveness of motile cilia-types within the vertebrate body, which inspires the need to evolve the cryo-EM technology so that, like the (9+2) cilia, the architecture of other kinds of cilia can also be analysed at atomic resolution.

## Results

### Cfap53 and Mns1 are paralogous MIPs

In order to establish whether Cfap53 and Mns1 are paralogous proteins we studied their evolutionary relationships. Initially, we utilized the MirrorTree software ^18^ to determine their phylogenetic relationship. We observed that 59 organisms were common among the generated trees, and a correlation value of 0.895 was obtained with a *p*-value significance of ≤ 0.000001. Therefore, these proteins exhibit a strong co-evolutionary relationship. Although, they share low sequence identity (22.8%) and similarity (48.9%), respectively (**Supplementary Figure 1**), it is highly likely that they are paralogs.

We also prepared a phylogenetic tree to better characterize the relationship between the two proteins. We found that Cfap53 and Mns1 co-cluster, and are possibly paralogous in *Ornithorhynchus anatinus*, *Sarcophilus harrisii*, *Monodelphis domestica*, *Rattus norvegicus*, and *Mus musculus* (**Figure 1E** and **Supplementary Table 1**).

### Mns1 localizes to the ciliary axoneme of (9+0) and (9+2) cilia in zebrafish and mice

Immunolocalization studies of endogenous or overexpressed tagged version of Mns1 in mouse sperm flagella, cilia in zebrafish LRO and human airways as well as cryo-EM analysis have suggested that Mns1 is an axonemal protein ^4,5,17,19,20^. Nevertheless, to investigate whether like Cfap53, Mns1 has any localization to additional ciliary regions that may have been overlooked in these earlier studies, we used a C-terminal haemagglutinin (HA) tagged Mns1 protein variant to visualize localization to cilia within the zebrafish KV and pronephric (kidney) ducts, that have (9+0) and (9+2) cilia-types, respectively. In both instances, we found that the localization of the Mns1-HA chimeric protein was restricted exclusively to ciliary axonemes (**Figure 2A,B**). Since the Mns1-HA protein also rescued the laterality defects of *mns1* zebrafish mutants (see below), we argue that this pattern represents the physiological localization of the protein.

**Figure 2.**
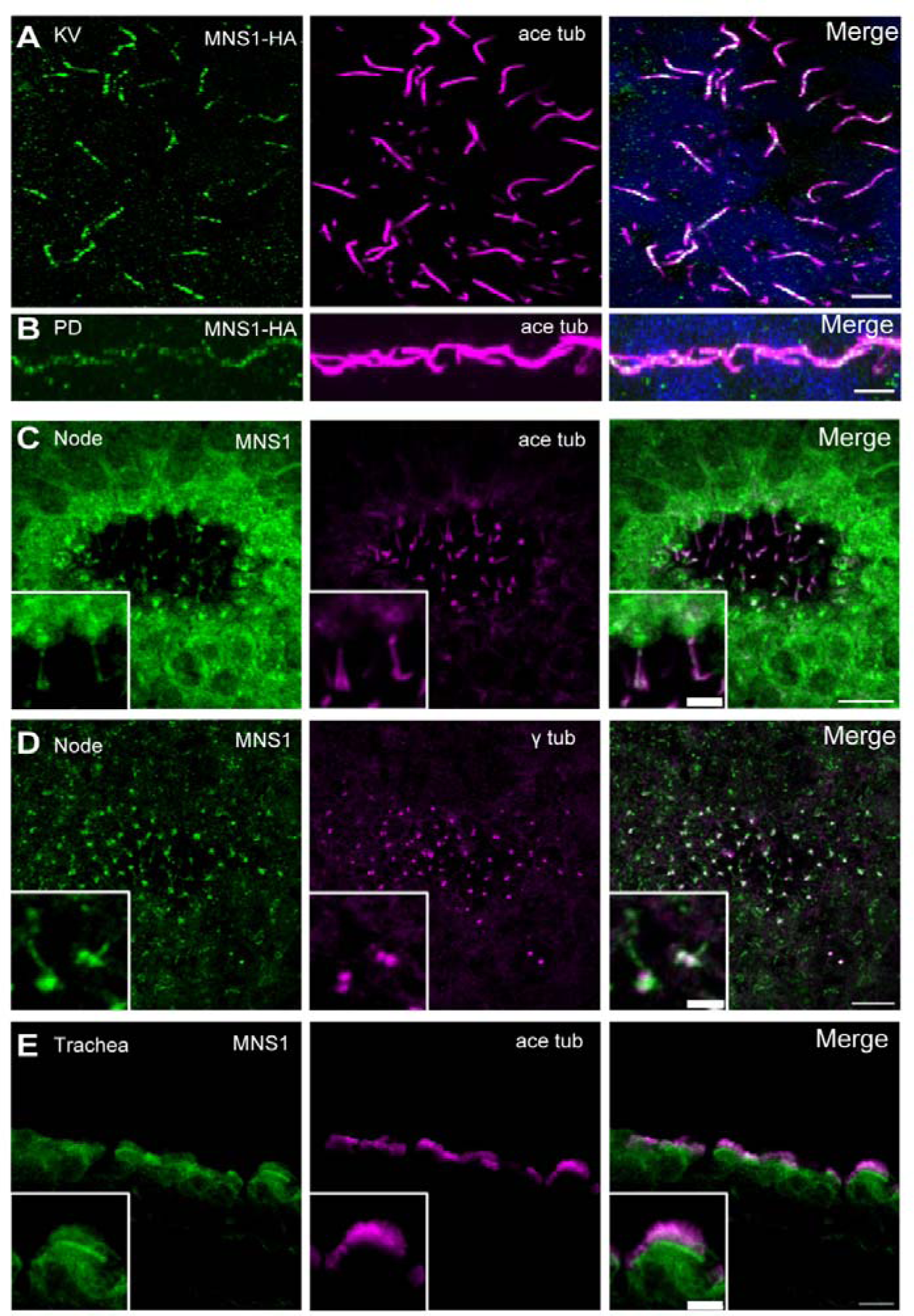
Localization of Mns1 to motile cilia in zebrafish and mouse tissues. (A) Immunofluorescence staining with antibodies to HA epitope (green) and acetylated tubulin (magenta) of cilia in zebrafish KV. (B) Immunofluorescence staining with antibodies to HA (green) and acetylated tubulin (magenta) of cilia in zebrafish pronephric duct (PD). (C) Immunofluorescence staining with antibodies to Mns1 (green) and acetylated tubulin (magenta) of cilia in the mouse node at E8.0. (D) Immunofluorescence staining with antibodies to Mns1 (green) and γ tubulin (magenta) of the mouse node at E8.0. Confocal gain in (C) was set higher than in (D) as Mns1 staining of the axoneme is weaker compared to the ciliary base. (E) Immunofluorescence staining with antibodies to Mns1 (green) and acetylated tubulin (magenta) of adult mouse tracheal cilia. Scale bars = 5 µm (A,B), 2 µm (C, D) and 5 µm (E). Insets show higher magnification views. Scale bars = 10 µm.

As mentioned before, in mouse embryos, motile (9+0) cilia are found in the central region of the node (the pit) at embryonic day 8 (E8) ^21^. Using an Mns1-specific antibody ^17^, we could localize the Mns1 protein to the axonemes of the node motile cilia (**Figure 2C**). Co-staining for Mns1 and γ−tubulin (**Figure 2D**) indicated that Mns1 is also localized to the base of the node cilia. However, this localization pattern is somewhat different from that of Cfap53, which is found mainly at the base of node cilia, the centriolar-satellites ^12^. In the mouse trachea, Mns1 localized along the axoneme and base of respiratory cilia (**Figure 2E**), similar to Cfap53 ^12^.

### Loss of Mns1 in zebrafish severely affected the motility of LRO cilia

To investigate the function of Mns1 in ciliary motility, we first used the zebrafish model to interrogate its role. Our earlier work with the Foxj1 transcription factor, a master regulator of motile cilia biogenesis, had identified *mns1* as one of its target genes ^19^. Consistent with this, published expression pattern of *msn1* has revealed that the gene is expressed in tissues that differentiate motile cilia in the zebrafish embryo (https://zfin.org/ZDB-GENE-030521-42/expression). We used the CRISPR-Cas9 gene editing technique to introduce a 19 bp deletion within exon 3 of *mns1* (**Figure 3A,B** ; see also Materials and Methods and **Supplementary Table 2**). Conceptual translation of the mutant cDNA predicted a highly truncated protein that is likely to be completely nonfunctional (**Figure 3C**). Despite this, we failed to observe canonical cilia defect-associated morphological anomalies among embryos obtained from crosses of heterozygous fish, such as ventral curvature of the body (that arises from ciliary defects within the brain ventricles and spinal canal) or cystic kidneys (triggered by defects in cilia within the kidney tubules). However, a significant proportion of the embryos (about 1/8, i.e half of the zygotic mutants) exhibited aberrant laterality of the heart, signifying defects of LRO cilia (**Figure 3D**). Since *mns1* mRNA is deposited maternally (**Supplementary Figure 2**), we reasoned that much stronger ciliary defect-associated phenotypes could possibly be apparent in maternal-zygotic (mz) mutants. Therefore, we raised the homozygous zygotic mutants to adulthood and obtained pure clutches of mz homozygous mutants through in-cross. In these embryos, we noted the same laterality defects of the heart, but now in a much higher proportion (approximately 1/2 of every clutch) (**Figure 3D**), and visualization of KV of these mz mutants confirmed strongly dysmotile cilia (**Supplementary Videos 1 and 2**). We could rescue the heart laterality defects using microinjection of sense mRNA encoding Mns1-HA into the mz mutant eggs (**Figure 3E**), confirming the specificity of the mutation and the physiologically relevant localization pattern of Mns1-HA to ciliary axonemes. However, like the zygotic mutants, the mz mutants did not exhibit axial curvature or kidney cysts, and videomicroscopy of (9+2) cilia of multiciliated cells (MCCs) of the nose showed a wild-type beating pattern (**Supplementary Videos 3 and 4**). These findings clarify that even though *mns1* mRNA is deposited maternally, it does not contribute to the masking of the ciliary defects in the zygotic mutants, and normal motility of (9+2) cilia and absence of phenotypic abnormalities, like axial defects and kidney cysts, could be due to redundancy with Cfap53.

**Figure 3.**
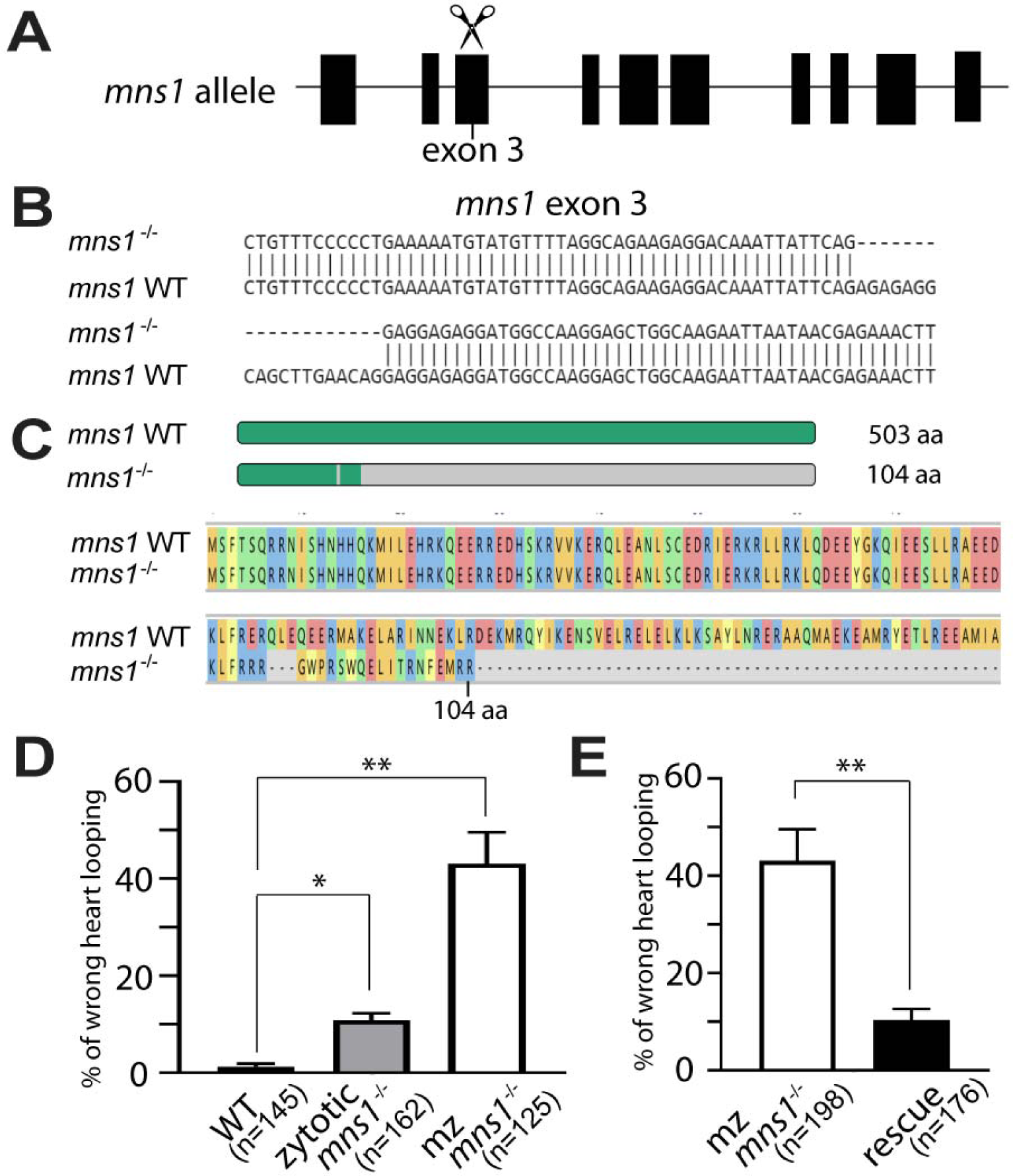
Generation of *mns1* mutant zebrafish and left-right asymmetry defects in the mutants. (A) Schematic diagram of zebrafish *mns1* locus and targeting of exon 3 with guide RNA. (B) Nucleotide sequence of the *mns1* mutant allele compared with the wild-type *mns1* gene. (C) Schematic and amino acid sequence of the predicted Mns1 mutant protein compared to the wild-type. (D) Percentage of embryos at 48 hpf showing heart looping defect compared with the wild-type embryos (sum of 3 technical replicates). (E) Percentage of *mns1* homozygous mutant embryos showing heart looping defect after rescue with wild-type *mns1-HA* mRNA compared with *mns1* homozygous mutant embryos at 48 hpf (sum of 3 technical replicates). Data are presented as mean ± SD, two tailed Student’s t-test (*p≤ 0.05, **p≤ 0.01).

### Mns1 knock out phenotype in the mouse

We also generated a mutant strain of mice lacking exon 3 of the *Mns1* gene (**Figure 4A,B** and **E**); the mutant protein is predicted to be severely truncated, and thus, likely to be nonfunctional (**Figure 4C,D**). About 30% of *Mns1* ^-/-^ mice died at the birth, and the about 40% of them succumbed within 12 weeks (**Figure 4F**). Mutant mice that perished between 30-60 weeks after the birth exhibited hydrocephalus (3/3 mice). When 12 *Mns1* ^-/-^ mice were examined for laterality defects in visceral organs either at birth or later, 9/12 showed laterality defects (**Table 1**). Two of them showed *situs inversus totalis* ; the remaining seven *Mns1* ^-/-^ mice examined showed a variable degree of heterotaxy. Abnormal lung lobation (9/12), reversed arching of the aorta (8/12), heart apex on the right (8/12), stomach on the right (6/11) were frequently observed (**Table 1**). Moreover, *Mns1* ^-/-^ males were infertile (2/2), while *Mns1* ^-/-^ females were fertile (2/2).

**Figure 4.**
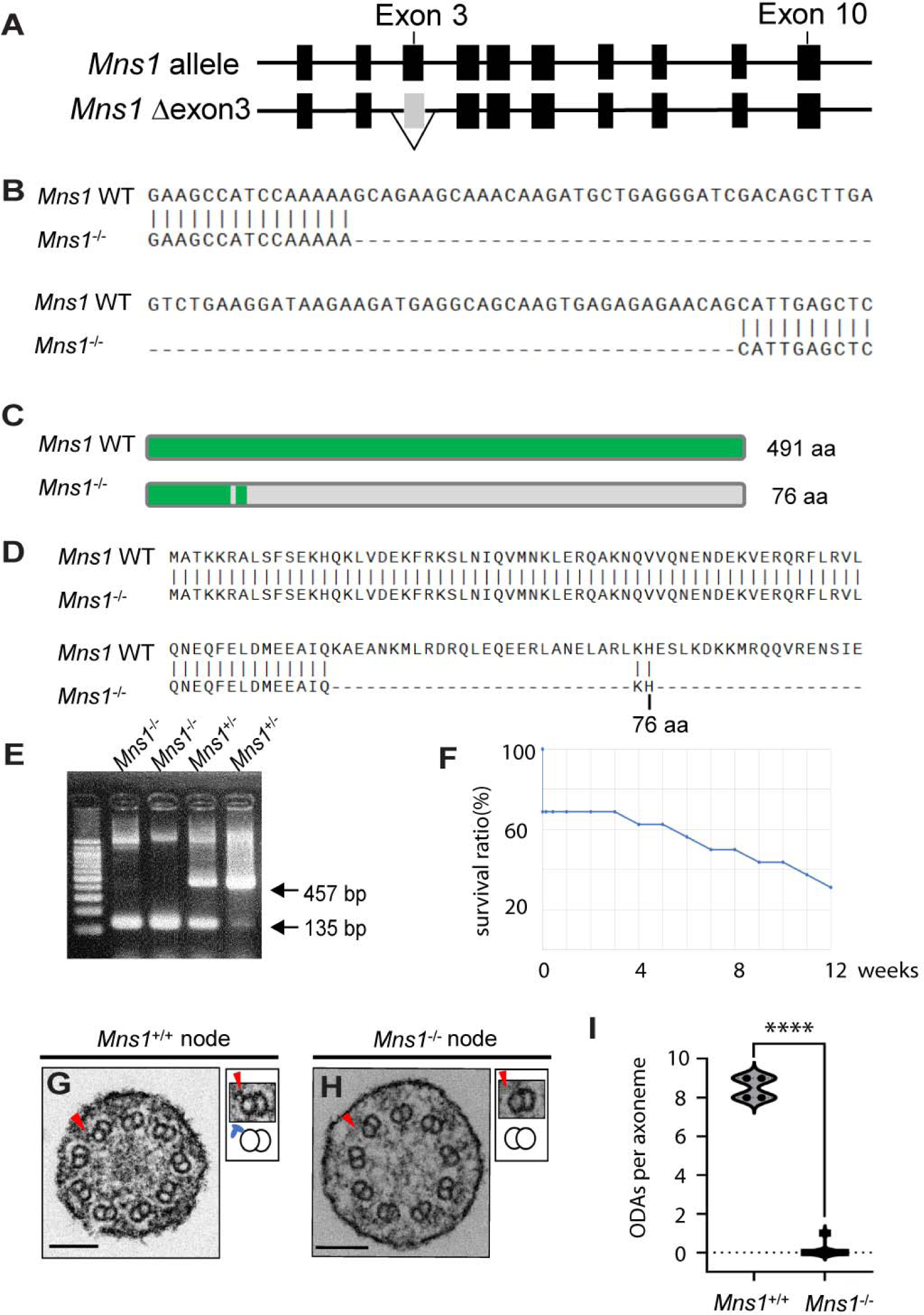
*Mns1* mutant mice lack ODAs from (9+0) nodal cilia. (A) Genetic structure of the wild-type mouse *Mns1* locus and the generation of knockout allele lacking exon 3. (B) Nucleotide sequence of the *Mns1* mutant allele compared with the wild-type *Mns1* gene. (C, D) Schematic (C) and amino acid sequence (D) of the predicted Mns1 mutant protein compared to the wild-type. (E) PCR genotyping of *Mns1* ^-/-^ mice. 457 base pair band (bp) represents the wild-type and 135 bp band represents the deletion allele, respectively. (F) Survival curve of *Mns1* ^-/-^ mice. (G, H) TEM data of wild-type and *Mns1* ^–/–^ node cilia. Higher magnification views of corresponding doublet microtubules indicated by the arrowheads, together with schematic diagrams of the microtubule pairs and ODAs (blue protrusions). Scale bars = 100 nm. (I) The number of ODAs per axoneme of node cilia from wild-type and *Mns1* ^-/-^ embryos. Data are presented as mean ± SD, two tailed Student’s t-test (****p= <0.0001).

**Table 1:** Abnormal left-right (L-R) phenotypic defects of *Mns1^-/-^* mice.

| L-R phenotypes observation | No. of abnormal L-R phenotype | Total observations | Incidence (%) |
| --- | --- | --- | --- |
| Heart apex on the right side | 7 | 11 | 63.6 |
| Abnormal lung lobation | 7 | 10 | 70.0 |
| Aortic arch on the right side | 8 | 11 | 72.7 |
| Azygos vein on the right side | 8 | 10 | 80.0 |
| Stomach on the right side | 6 | 8 | 75.0 |
| Abnormal liver lobation | 7 | 8 | 87.5 |
| Vena cava located to the left of the aorta | 8 | 9 | 88.9 |

As suggested by the laterality defects, node cilia of *Mns1* ^-/-^ embryos were almost completely immotile (**Supplementary videos 5 and 6**). Analysis of the structure of these cilia by transmission electron microscopy (TEM) showed a complete absence of ODAs from their axonemes (**Figure 4G-I**). Tracheal cilia of *Mns1* ^-/-^ mice were motile, but showed abnormal beating pattern (**Supplementary videos 7 and 8**). In particular, the beating angle of the tracheal cilia was smaller, rendering the effective stroke inefficient, similar to what we have previously reported for *Cfap53* mutants (for TEM data on tracheal cilia of *Mns1* mutants, see subsequent section).

### Concomitant loss of Mns1 and Cfap53 severely affected motility of (9+2) cilia in the zebrafish

Our previous work with antisense morpholinos and the study of Noel et al. with a *cfap53* mutant strain of zebrafish had established that the protein is essential for motility of KV cilia ^11,14^. To examine potential redundant roles of Cfap53 and Mns1, we generated an independent allele of *cfap53* using CRISPR/Cas9 gene editing (**Figure 5A**). This mutant allele consists of a 14 bp deletion in exon 2 (**Figure 5B,C** ; the allele described in Noel et al. ^14^ is a 7 bp deletion within the same exon). *cfap53* mRNA is not deposited maternally (**Supplementary Figure 2**), and we raised the embryos obtained from het incrosses to adulthood. Incross of the homozygous mutant fishes yielded mz mutants which did not exhibit any other ciliary abnormalities besides KV cilia motility defects (**Supplementary video 9**) and abnormal left-right asymmetry (**Figure 5D**), recapitulating the severe loss of KV cilia motility and laterality defects reported by Noel et al. as well as in our study with the morpholino ^11,14^. We also examined the motility of (9+2) MCC cilia of the nose and could not discern any obvious deviations from the wild-type (**Supplementary video 10**). Thus, like Mns1, Cfap53 is dispensable for (9+2) cilia motility in the zebrafish, pointing to a likely redundancy with Mns1.

**Figure 5.**
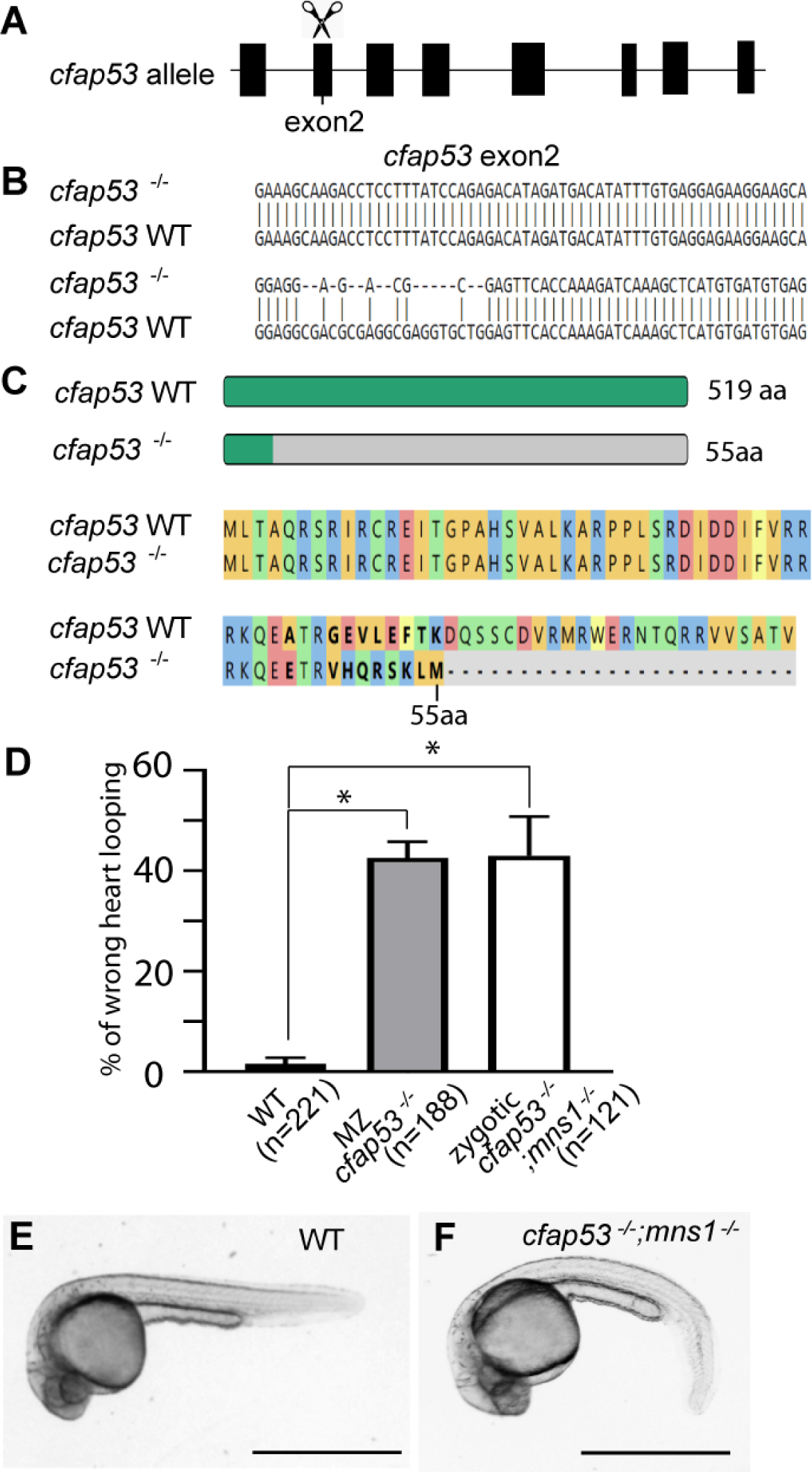
Zebrafish *cfap53* and *mns1* mutants exhibited severe ciliary disorder phenotypes. (A) Schematic diagram of zebrafish *cfap53* locus and targeting exon 2 with guide RNA. (B) Nucleotide sequence of the *cfap53* mutant allele compared with the wild-type gene. (C) Schematic and amino acid sequence of the predicted Cfap53 mutant protein compared to the wild-type. (D) Percentage of embryos showing heart looping defect at 48 hpf compared with wild-type control embryos (sum of 3 technical replicates). Data are presented as mean ± SD, two tailed Student’s t-test (*p≤0.05). (E) A wild-type zebrafish embryo morphology at 24hpf. (F) A *cfap53*^-/-^; *mns1*^-/-^ double mutant embryo with curved body axis. Scale bar = 1mm.

To investigate the possibility of redundancy, we used this allele of *cfap53* to generate double heterozygous *mns1*^+/-^; *cfap53*^+/-^ fish, and in-crossed them to obtain zygotic double mutants for *mns1* and *cfap53*. As expected from segregation based on Mendelian genetics, we found that approximately 1/16 of the embryos from such crosses exhibited a prominent axial curvature (**Figure 5E,F**), typical of zebrafish with strong cilia motility defects, that was not apparent in the single mutants for either of the genes. Genotyping these ventrally curved embryos confirmed them to be double mutants. The double mutants also exhibited heart laterality defects, the hall mark phenotype of the single mutants, with about 50% of the double mutants having the heart tube looped on the wrong side (**Figure 5D**). Despite the prominent axial curvature defect, the double mutants did not develop cystic kidneys suggesting that the motility of (9+2) cilia may not be fully compromised even with the combined loss of both the MIPs. Consistent with this notion, when we analysed (9+2) cilia of the MCCs in the nose, a range of motility defects was apparent. In all double mutant embryos examined, these cilia exhibited aberrant motion distinct from the smooth wave form characteristic of wild-type embryos (**Supplementary video 11**). In a proportion of the double mutants (about 20%), we also found complete paralysis of the majority of the cilia, and those that were still motile, showed uncoordinated twitching-like activity (**Supplementary video 12**). Since the *mns1*; *cfap53* zygotic homozygotes are embryonic lethal due to their curved body axis precluding acquisition of swimming ability, we generated *mns1*^-/-^; *cfap53*^+/-^ adults, and bred them to obtain mz mutant *mns1* and zygotic mutant *cfap53* double homozygotes. These mutants exhibited the same spectrum of phenotypic defects and (9+2) nasal cilia abnormalities as the zygotic *mns1*; *cfap53* double mutants (data not shown). Since KV cilia were immotile or exhibited strong motility defects in the *cfap53* and *mns1* single mutants, respectively, we did not analyse motility of these cilia in the double mutants.

### (9+2) cilia of *Mns1* ; *Cfap53* double mutant mice are largely devoid of ODAs

Genetic interaction between *Mns1* and *Cfap53* was also apparent in the mouse. First, the double mutant mice, lacking both Mns1 and Cfap53 function, showed a much higher incidence of lethality after birth. Unlike *Mns1* and *Cfap53* single mutant mice, about 60% of the double mutants died at birth, and most of them perished within 5 weeks after birth (**Figure 6A**). Secondly, the double mutants exhibited more severe laterality defects than *Mns1* or *Cfap53* single mutant mice. In particular, the incidence of azygous vein on the right, stomach on the right and abnormal liver lobation was higher in the double mutants (**Figure 6B,C** and **Table 2**). Finally, tracheal cilia of *Mns1* ; *Cfap53* double mutant mice lost ODAs almost completely, while the loss of ODAs in tracheal cilia was only partial in the single mutants for each gene (**Figure 6D-G** ; see also ^12^). Due to the early lethality of the double mutants, we were unable to perform videomicroscopy to analyse ciliary beating as the few trachea that we could isolate from these animals was used for TEM analysis.

**Figure 6.**
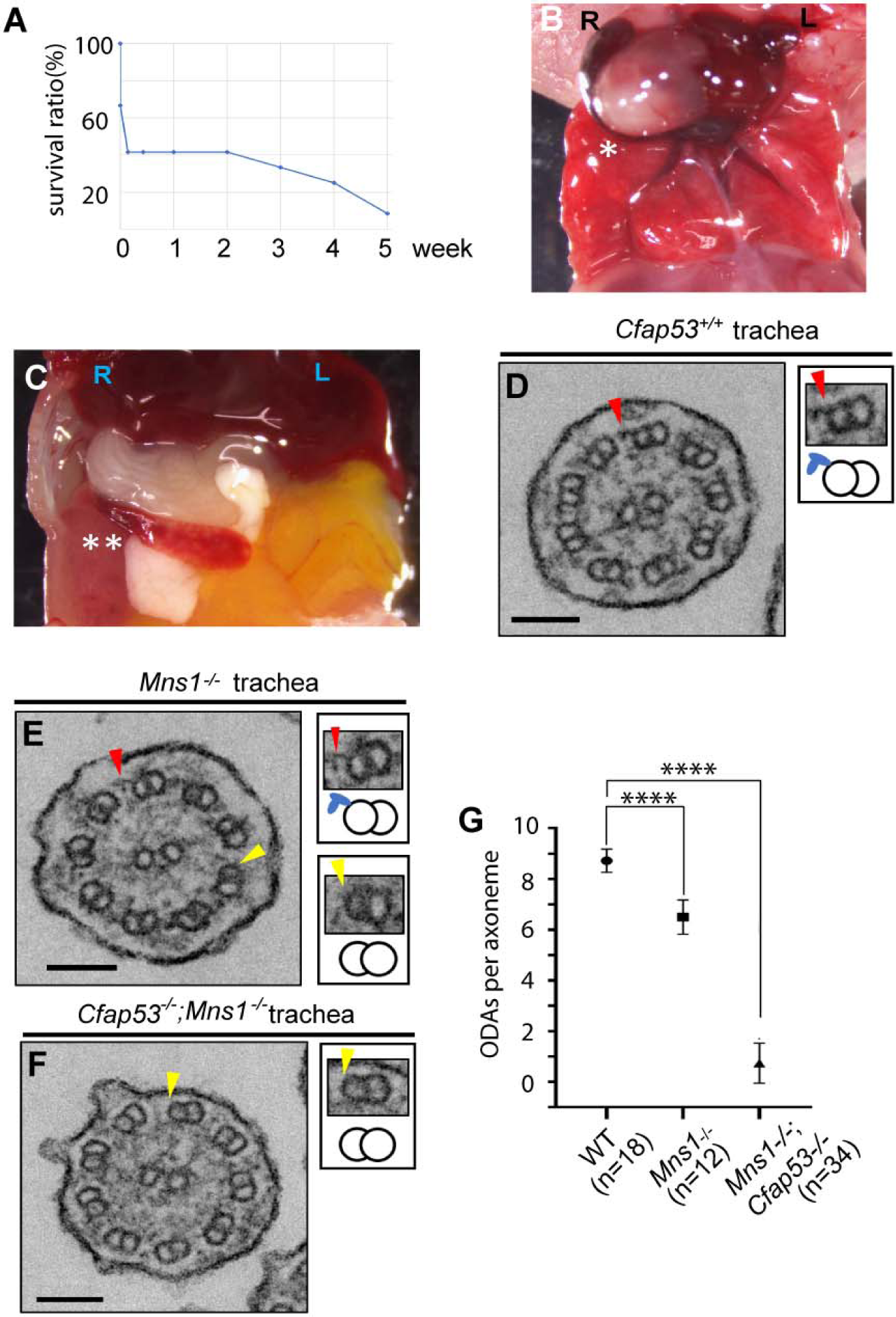
Loss of ODAs from (9+2) tracheal cilia of *Cfap53* ; *Mns1* double mutant mice. (A) Survival curve of *Cfap53*^-/-^; *Mns1* ^-/-^ double mutant mice. (B, C) Laterality defects of *Cfap53*^–/–^, *Mns1* ^-/-^ mice. Asterisk indicates heart apex in the reversed position (B) and double asterisk indicates spleen in the reversed position (C). (D-F) TEM data of tracheal cilia are represented as *Cfap53*^+/+^ (D), *Mns1* ^–/–^ (E) and *Cfap53*^–/–,^ *Mns1* ^-/-^ (F). Insets show higher magnification views of corresponding doublet microtubules indicated by the arrowheads, together with schematic diagrams of the microtubule pairs and ODAs (blue protrusions). Scale bars = 100 nm. (G) The number of ODAs per axoneme of tracheal cilia from wild-type control, *Mns1* ^-/-^ and *Mns1* ^-/-^; *Cfap53*^-/-^ adult mice. Data are presented as mean ± SD, two tailed Student’s t-test (****p= <0.0001).

**Table 2:** Abnormal L-R phenotypic defects of *Mns1*^-/-^; *Cfap53*^-/-^ mice.

| L-R phenotypes observation | No. of abnormal L-R phenotype | Total observations | Incidence (%) |
| --- | --- | --- | --- |
| Heart apex on the right side | 8 | 12 | 66.7 |
| Abnormal lung lobation | 9 | 12 | 75.0 |
| Aortic arch on the right side | 8 | 12 | 66.7 |
| Azygos vein on the right side | 3 | 7 | 42.9 |
| Stomach on the right side | 6 | 11 | 54.5 |
| Abnormal liver lobation | 6 | 9 | 66.7 |
| Vena cava located to the left of the aorta | 7 | 9 | 77.8 |

### Cfap53 interacts with Mns1, but unlike Mns1 does not self-associate

In our earlier work with Cfap53, we provided evidence that the protein can interact with the ODA docking complex member Odad4 as well as several ODA dynein proteins in immunoprecipitation assays ^12^. In addition, based on the localization of Cfap53 to centriolar satellites of the basal bodies and the established role of the satellites in facilitating protein transport into the axoneme, we proposed that Cfap53 could also be involved in the transport of Odad4 and ODA dyneins into cilia ^12^. In line with these interactions, the ODA complex as well as Odad4 was lost from nodal cilia of Cfap53 mutant mice, but in (9+2) tracheal cilia, there was only a partial loss of the ODAs while Odad4 localization remained unaffected ^12^. Biochemical studies with Mns1 have also shown that it can interact with another ODA docking complex member, Odad1 (aka Ccdc114) ^17^. Additionally, Mns1 has been shown to interact with itself, forming dimers and oligomers ^20^. Indeed, end-to-end self-association for Mns1 as well as Cfap53 has been noted in the cryo-EM studies, allowing them to generate a 48 nm periodicity along the length of the axoneme (**Figure 1D**) ^4,5^. Moreover, the atomic models from cryo-EM studies predict interaction of Mns1 with Cfap53 as well (**Figure 1D**) ^4,5^. To validate Cfap53 self-association and interaction of Cfap53 with Mns1, we performed immunoprecipitation studies with over-expressed epitope tagged proteins in cultured HEK293T cells. Using this assay, we observed that CFAP53-Myc can immunoprecipitate with MNS1-HA (**Figure 7A**). However, unlike MNS1, we were unable to detect any self-association between CFAP53-Myc and CFAP53-HA (**Figure 7B**). Consistent with this, while MNS1-HA and MNS1-Myc formed oligomeric filaments in the cytoplasm of HEK293T cells as reported previously ^20^, we failed to observe such filaments with CFAP53 (**Figure 7C,D** ; also see Discussion).

**Figure 7:**
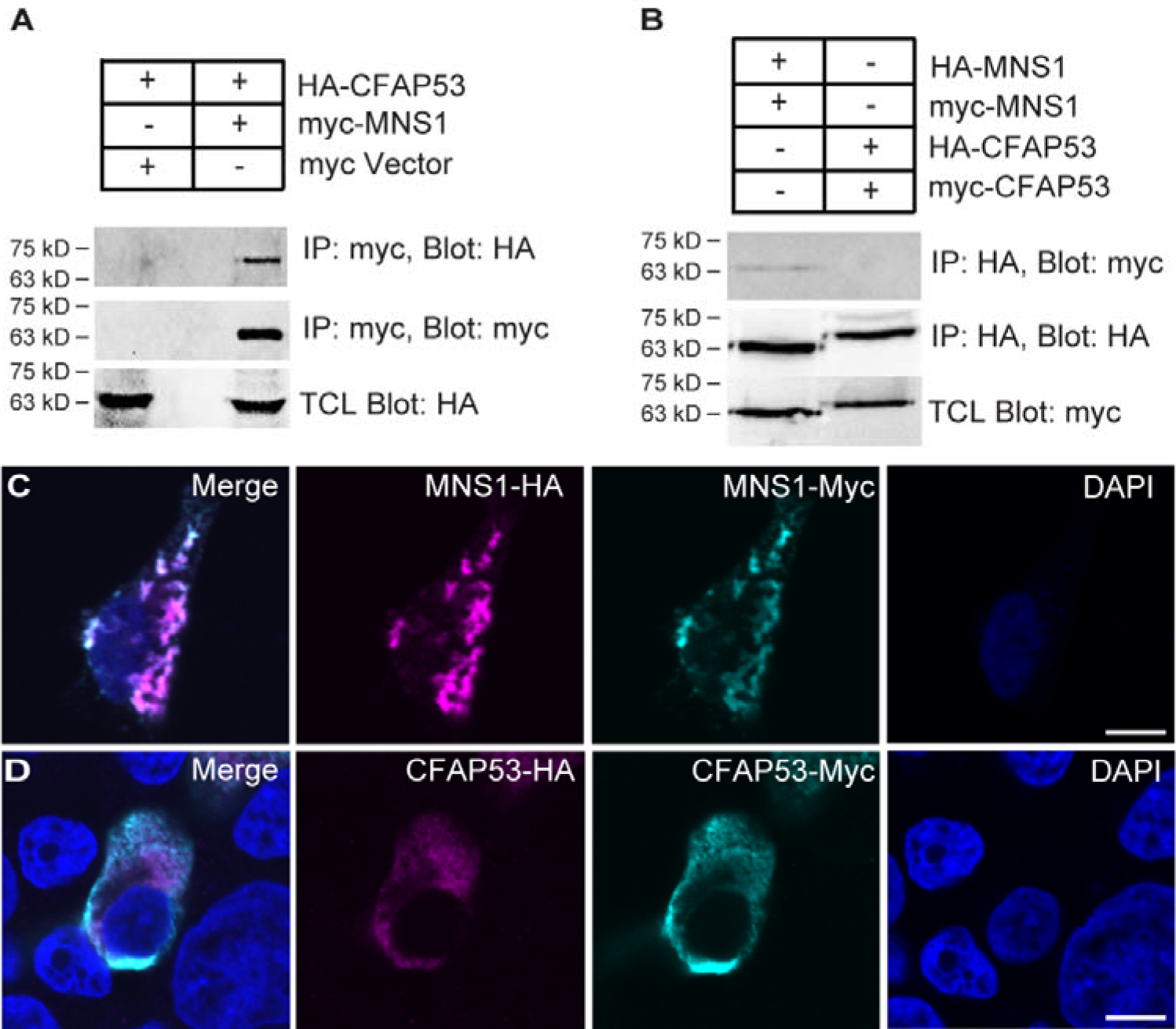
Biochemical interaction between Mns1 and Cfap53 proteins. (A) Myc tagged MNS1 immunoprecipitated with HA tagged CFAP53. MNS1-Myc and CFAP53-HA plasmids were co-transfected into HEK293T cells. CFAP53-HA was pulled down by immunoprecipitation of MNS1-Myc. (B) MNS1 but not CFAP53 exhibited self-interaction. CFAP53-HA and CFAP53-Myc and MNS1-HA and MNS1-Myc were co-transfected into HEK293T cells. MNS1-Myc was pulled down by immunoprecipitation of MNS1-HA. However, CFAP53-Myc was not pulled down by immunoprecipitation of CFAP53-HA. (C) MNS1 formed filament-like structures in HEK293 cells, after co-transfection of MNS1-Myc and MNS1-HA expression plasmids. MNS1-HA was stained with anti-HA antibody (magenta), while MNS1-Myc was stained with anti-Myc antibody (cyan). (D) CFAP53 showed diffuse staining pattern in HEK293 cells after co-transfection of CFAP53-Myc and CFAP53-HA expression plasmids. CFAP53-HA was stained with anti-HA antibody (magenta), while CFAP53-Myc was stained with anti-Myc antibody (Cyan). Scale bar = 5 µm. All interactions and localization studies were performed in 2 technical replicates.

### Distinct localization of ODA-DC members to (9+0) versus (9+2) cilia in zebrafish

Besides Cfap53 and Mns1, which we have established to have distinct localization and/or function in (9+0) versus the (9+2) cilia, we have previously shown that in the mouse, individual ODA dyneins also have unique patterns of expression, localization and function in these two cilia-types. Thus, in mice, while Dnah11 is localized along the entire length of the axonemes of LRO cilia, it is localized specifically to the proximal region of tracheal cilia ^12^. Even more intriguingly, the Dnah9 protein, which localizes to the distal axoneme of tracheal cilia, is not expressed in the node despite the gene being transcribed in node cells ^12^. These observations illustrate that besides the traditional view that the absence of the central pair and associated structures like the radial spokes is the key defining difference between (9+0) and (9+2) cilia, several other ciliary components have differential expression, localization and/or function in the two cilia-types. Since cryo-EM level ultrastructural studies with (9+0) cilia are currently not possible, we decided to further explore the differences between (9+0) and (9+2) cilia by evaluating localization patterns of a set of evolutionarily conserved and key motility components – the ODA-DC members Odad1, -3 and -4. We used N-terminal myc tag for all 3 proteins, and for the C terminus we used GFP for Odad1 and -3, whereas for Odad4 we used the HA tag, respectively. We injected the synthetic sense mRNAs encoding each of the N- or C-terminally tagged proteins into zebrafish eggs and analysed their localization to cilia in KV, kidney ducts and spinal canal; TEM analysis of the latter has revealed them to be largely of the (9+0) type ^22,23^. For Odad1 and -3 proteins, N-terminally myc tagged versions localized almost exclusively to the axonemes of the (9+2) kidney duct cilia, consistent with reports of localization of the endogenous proteins with specific antibodies to (9+2) cilia of mammals ^24–26^ (**Figure 8A,B**); however, we failed to observe any localization of myc-Odad4 (data not shown). Similarly, C-terminal GFP tagged Odad1 and -3 as well as C-terminal HA tagged Odad4 localised along the axonemes of kidney duct cilia; there was also substantial localization of Odad4-HA in the cytoplasm (**Figure 8C** and **Supplementary Figure 3**). By contrast, N-as well as C-terminally tagged Odad1 and -3 proteins localised exclusively to the base of all KV well as spinal canal cilia that we analysed, whereas for Odad4, localization of HA-tagged C-terminal version was observed along their axonemes, similar to what we have reported previously for (9+0) LRO cilia of the mouse (**Figure 8D-I** and **Supplementary Figure 3**); there was also substantial Odad4-HA accumulation in the cytoplasm, and we again failed to observe localization of myc-Odad4 to these cilia (data not shown; see Discussion on why N-terminal tag could interfere with Odad4 localization).

**Figure 8:**
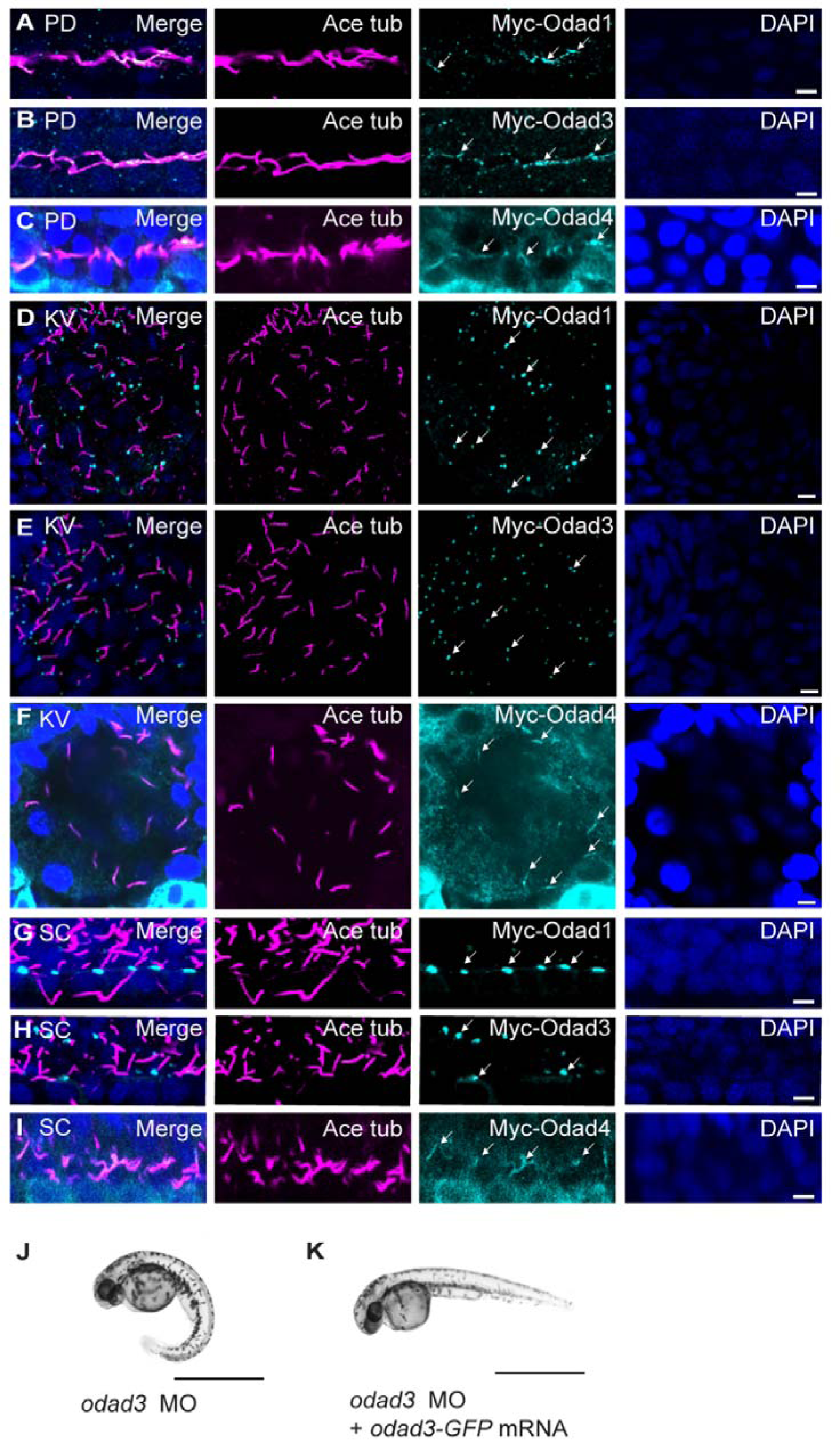
Differential localization of Odad proteins to the axoneme and basal bodies of zebrafish motile cilia. (A-C) Odad1, -3 and -4 localized on the axoneme of (9+2) cilia in the zebrafish pronephric duct (PD) at 24 hpf. (D, E) Odad1 and -3 localized at the base of KV cilia at the 10 somites stage. (F) Odad4 localized on the axoneme of KV cilia. (G, H) Odad1 and -3 localized at the base spine canal cilia at 24 hpf. (I) Odad4 localized to the axoneme of spine canal (SC) cilia. The tagged Odad proteins were detected by staining with anti-myc antibody (for Odad1 and -3; cyan) or anti-HA antibody (for Odad4; cyan) (indicated by arrows), cilia were stained with anti-acetylated tubulin antibody (magenta) and nuclei with DAPI. Scale bar = 5 µm. (J) An *odad3* morphant showing ventrally curved body axis (95% morphants exhibited curved body axis). (K) An *odad3* morphant (MO) with rescue of axial curvature defect on co-injection with *odad3-gfp* mRNA (90% of morphants rescued to wild-type). Scale bar = 1 mm. All localization, morpholino and rescue experiments were performed in 3 technical replicates.

To adduce evidence that these localization patterns of the tagged and over-expressed Odad proteins indeed reflect their endogenous localization patterns and do not compromise their function, we attempted to rescue *odad3* morphants with the N-terminal myc and C-terminal GFP tagged proteins. As described previously, *odad3* morphants exhibit ventrally curved body axis and defects in looping of the heart tube ^27^, closely mimicking *odad3* genetic mutants ^24^, and we were able to reproduce these phenotypes after injecting the morpholino into wild-type zebrafish eggs (**Figure 8J**). Analysis of KV cilia motility of the morphants revealed immotile or highly dysmotile cilia, in line with the overt laterality defects of the heart (**Supplementary video 13**). Notably, mRNAs for both the N-as well as the C-terminally tagged Odad3 proteins were able to significantly rescue KV cilia motility, axial curvature and heart *situs* defects of the morphants when co-injected with the morpholino, confirming that the ciliary localization patterns exhibited by the chimeric Odad3 proteins is physiological, and the epitope tags do not disrupt their activity (**Figure 8K** and **Supplementary video 14**).

## Discussion

We and others first identified Cfap53 as a putative ciliary protein based on transcriptomic and proteomic screens aimed at discovering novel ciliary/centriolar components ^19,28–31^. Subsequently, using antisense morpholino based loss-of-function studies with the zebrafish as well as genetic analysis of a patient with *situs inversus*, we had established that inactivation of Cfap53 is associated with the loss of ODAs from LRO cilia, rendering them immotile ^11^. These findings were independently validated by other groups, thereby establishing that Cfap53 is a key protein responsible for proper motility of LRO cilia and the establishment of asymmetry of visceral organs ^13–15^. Furthermore, investigating the localization pattern of Cfap53 led us and others to conclude that the protein has two distinct patterns of subcellular distribution: it is present on the ciliary axonemes and is also associated with the ciliary basal bodies as a likely constituent of centriolar-satellites ^11,13^. More recently, we used genetic and cell biological analysis in the mouse to further investigate the localization and function of Cfap53 ^12^. Consistent with the zebrafish data, we could establish that Cfap53 is also a motile cilia protein that exhibits differential localization in (9+0) versus the (9+2) cilia, being present mainly at the basal bodies of the former and at the basal bodies as well as along the axonemes of the latter. Moreover, loss of Cfap53 preferentially affected the motility of the (9+0) cilia, with the (9+2) cilia only being altered marginally in their beat frequency and amplitude. Why Cfap53 has this differential requirement in the two cilia-types has remained an outstanding question. On the other hand, the function of Mns1, another filamentous MIP, was originally investigated based on its prominent expression in the mouse testes, specifically in the spermatocytes, spermatids and along the axonemes of mature sperm ^20^. Consistent with these data, mice mutant for *Mns1* developed sperm with rudimentary flagella that were highly disorganised at the ultrastructural level. In addition, the mutant mice also exhibited randomized left-right asymmetry, hydrocephalus and TEM analysis of their tracheal (9+2) cilia showed a partial loss of the ODAs ^20^. However, motility defects, if any, of (9+0) or the (9+2) cilia were not investigated in that study. Like Cfap53, Mns1 has also been associated with *situs* abnormalities and male infertility in humans ^16,17^. Moreover, like Cfap53, which can interact with the ODA dyneins as well as the ODA-DC member Odad4, Mns1 has been shown to associate with the ODA-DC protein Odad1 ^17^.

It is, however, cryo-EM based structural analyses of *Chlamydomonas* flagella and mammalian respiratory cilia, both with a (9+2) configuration, which have revealed that in the ciliary axonemes, Cfap53 and Mns1 are filamentous MIPs which decorate the lumens of the A microtubules of the axonemal doublets ^4,5^. Cfap53 and Mns1 occupy clefts between adjacent microtubule protofilaments – Cfap53 between protofilaments A6 and A7 and Mns1 between A7 and A8. In view of this neighbouring localization pattern, their interactions with ODA-DC components and loss of the ODAs when the proteins are mutated led us to investigate if they have redundant roles in regulating the motility of the (9+2) cilia. We have now established that Cfap53 and Mns1 are paralogous proteins and consistent with this, in mice and zebrafish doubly mutant for both proteins, motility of the (9+2) cilia is strongly compromised and the ODAs are almost completely lost.

In addition to analysis of the double mutants, we have performed detailed analysis of cilia structure and motility in the *Mns1* single mutants as well, that have not been examined in the earlier report ^20^. While the motility of LRO cilia in zebrafish as well as mice was strongly affected, in both organisms (9+2) cilia retained motility, albeit with partial loss of the ODAs. These data collectively establish that Cfap53 and Mns1 are both essential for LRO cilia motility, but their individual functions are largely dispensable in the (9+2) cilia. We propose that this disparity likely arises from their distinct localization patterns and activities in the two kinds of cilia. In the (9+0) LRO cilia, Cfap53 possibly facilitates transport of ODA dyneins and ODA-DC members as part of the centriolar-satellites, while Mns1 functions as a MIP for stable anchoring of the ODA-DC and ODAs. On the other hand, in (9+2) cilia, both proteins predominantly function as MIPs for ODA-DC and ODA attachment to the axonemal microtubules, thus, resulting in a considerable degree of redundancy. An intriguing observation that has emerged from our study of the zebrafish *mns1* and *cfap53* mutants is that even though cilia within the spinal canal, which drive CSF flow and formation of the glycoproteinaceous Reissner fibre for the development of a straight body axis ^32^, have been described to be (9+0) ^22,23^, no axial defects were apparent in the single mutants during embryonic, larval or adult stages. This suggests that the (9+0) cilia of KV are distinct from those of the spinal canal with respect to their requirement for Mns1 or Cfap53 function. Likewise, even though we have organised our narrative based on KV cilia being (9+0), in reality they are a mixture of (9+0) and (9+2) ^5,33^ (the exact proportions of the two kinds of cilia in KV is presently not clear). Yet, most KV cilia exhibit rotational motion (apparent in **Supplementary video 1**), ciliary components like Cfap53, localizes exclusively to the base of all KV cilia and mutation of *cfap53* and *mns1* uniformly affects their motility, implying that KV (9+2) cilia could be distinct from those is other tissues, such as the pronephric ducts and the nose. Thus, the conventional (9+0) and (9+2) labels are at best a very crude way of classifying motile cilia, and our current study categorically establishes that these organelles in different tissues of the vertebrate body are considerably more diverse than presently recognized.

The cryo-EM data have also revealed that both Cfap53 and Mns1 associate in a head to tail fashion, forming a 48 nm repeat long the axoneme ^4,5^. For Mns1, over-expression in cultured cells as well as biochemical studies such as GST pull down, yeast-two hybrid analysis and immunoprecipitation have confirmed oligomerization (^20^ and our present study). By contrast, we were unable to demonstrate self-association of Cfap53 using immunoprecipitation of over-expressed proteins. However, conditions for co-immunoprecipitation reactions may not be congenial for preserving interactions that occur in the native condition. In view of this caveat, Cfap53 self-association is perfectly plausible in the context of the axonemal microtubular environment. In addition to self-association, the cryo-EM data also indicate interaction between the two proteins themselves, which we have been able to validate using immunoprecipitation. Furthermore, each protein has been proposed to associate with a complement of various other MIPs that is unique to each protein: These include Cfap21, C1orf158 and Cfap161 for Mns1 and Pierce1, Pierce2, and NME7 for Cfap53 ^4,5^. We have shown that Pierce1/2 proteins are necessary for ODA attachment (Gui, Farley et al. 2021): The validity and significance of the other associations will require further biochemical and genetic studies.

Similar to Cfap53 and Mns1, loss of function of a number of other MIPs, like Cfap45, Cfap52, Enkur and Nme7, have been documented to manifest in strong left-right asymmetry defects (signifying dysmotility of the LRO cilia) but relatively mild to indiscernible effect on the respiratory (9+2) cilia ^34–37^. Although it is not presently clear why these proteins have such differential requirements in the motility of the two kinds of cilia, none-the-less, the data provide additional support for our view that the (9+0) and (9+2) cilia have key structural differences. In the final section of our manuscript, we attempted to further probe the molecular differences between the (9+0) and (9+2) cilia using the zebrafish embryo. Rather surprisingly, we found that even highly evolutionarily conserved and core ODA-DC members localized differentially to KV, spinal canal and kidney tubule cilia. Thus, like Cfap53, Odad1 and -3 localized exclusively to the base of KV and spinal canal cilia (consistent with previous observation of Odad3 localizing to the base of spinal canal cilia ^27^), raising the intriguing possibility that in addition to their role in anchoring dynein arms to axonemal microtubules, these proteins could have other functions at the ciliary base. This could be transport of the dynein arms, as we have invoked for Cfap53, or other roles that are presently not defined. Exploring the interactome of these Odad proteins could provide valuable clues for these additional functions. Also, the absence of Odad1 and 3, central components of the pentameric vertebrate ODA-DC, from the axonemes of KV and spinal canal cilia suggests a significantly simplified ODA-DC or substitution of their function by other proteins in these cilia-types. For Odad4, we failed to observe any ciliary localization with the N-terminal myc tag. The first 10 amino acids at the N-terminus of Odad4 are not resolved in the cryo-EM structure, indicating that this region is flexible, and an N-terminal tag should not interfere Odad4 binding to microtubules. One possibility is that the N-terminal myc tag affected the transport of Odad4 into cilia. Another possibility is that the anti-myc antibody could have no access to the N-terminal tag because the N-terminus of Odad4 is located between the axonemal microtubules and ODA motor domain and this space is rather small (M. Gui, personal communication). Unfortunately, we were not able to determine whether Odad1 and -3 also localize to the base of (9+0) cilia of the mouse ventral node due to the lack of specific antibodies or reporter transgenes for the mouse proteins.

In conclusion, our work establishes that while ultrastructural data of (9+2) cilia have been key to our understanding of their molecular composition and mechanisms of motility, such information cannot be directly transposed to (9+0) cilia, and potentially to a variety of other motile cilia sub-types. Till we are able to obtain high resolution structural data for different kinds of cilia, genetics and cell biological investigations, as illustrated with this work, will continue to remain instrumental in enabling us to dissect their molecular composition and mechanisms of motility. How such a diversity of cilia-types is assembled in different tissues, and in some instances even within the same tissue, promises to be an enterprising area for future investigations.

## Acknowledgements

We thank H. L. Yeo for technical assistance with generation of the zebrafish *cfap53* mutant strain, B. Durand for plasmids containing zebrafish *odad1* and -*3* cDNAs and A. Brown and M. Gui for critical reading and invaluable comments on the manuscript. This work was supported by funds from the Ministry of Education, Culture, Sports, Science, and Technology (MEXT) of Japan (no. 17H01435) to H. H., the Council for Scientific and Industrial Research of India to S. C. and the Agency for Science, Technology and Research (A*STAR) of Singapore to S. R.

## Author contributions

SR and HH conceived the study; LH, VPK and KXC performed all of the experiments with zebrafish and protein biochemistry; WKT and YI performed all of the experiments with mice, IM did the bioinformatics analysis, SR and HH acquired funding and supervised the zebrafish and mouse experiments, respectively; SC supervised the bioinformatics analysis; SR wrote the paper with input from HH and SC; all authors read and endorsed the final version.

## Materials and Methods

### Protein ortholog or paralog identification and phylogenetic analysis

Protein sequences of human MNS1 and CFAP53 were obtained from the UniProt database ^38^. MNS1 and CFAP53 were aligned using EMBOSS needle ^39,40^ for the computation of sequence identities between them. Orthologs or similar sequences were determined using domain enhanced lookup time accelerated BLAST (DELTA-BLAST) ^41^. Sequences sharing an E-value of ≤ 1e-04, query length coverage of ≥ 70% and sequence identity of ≥ 45% with the query proteins from the NCBI non-redundant database ^42^ were taken as close orthologs. MNS1 and its orthologs (244 sequences) and CFAP53 and its orthologs (87 sequences) were aligned with an iterative refinement method (MAFFT-L-INS-I version 7) giving preference to local alignment such that the conserved regions are well aligned ^43,44^. Further, multiple sequence alignment (MSA) of orthologous proteins of MNS1 and CFAP53 were analysed with the help of the MirrorTree program ^18^. Organisms in which Mns1 and Cfap53 shared an evolutionary relationship were selected and a combined MSA and phylogenetic tree were prepared to determine whether these genes could be paralogous. Phylogenetic tree was generated in Randomized Axelerated Maximum Likelihood (RAxML version 8.1.6) ^45^ program that utilizes the maximum likelihood algorithm. The phylogenetic trees were visualized in FigTree (http://tree.bio.ed.ac.uk/software/figtree/;). Additionally, selected Mns1 and Cfap53 protein and gene sequences were aligned with the help of EMBOSS matcher ^40,45^.

### Zebrafish strains

All zebrafish strains were maintained according to established protocols for fish husbandry at the Institute of Molecular and Cell Biology (IMCB) zebrafish facility. The following wild-type and mutant strains were used in this study: AB (inbred wild-type control), *mns1* mutant (19 base pair (bp) deletion in the third coding exon), *cfap53* mutant (14 bp deletion in the second coding exon) (generated using CRISPR/Cas9 technology; for guide RNA sequences and genotyping primers see **Supplementary Table 2**) and *mns1*; *cfap53* double mutants. All experiments with the zebrafish were approved by the Singapore National Advisory Committee on Laboratory Animal Research (protocol number: 221702).

### Generation of *cfap53*^-/-^*, mns1* ^-/-^ and *cfap53* ^-/-^*; mns1*^-/-^ double mutant fishes

Single guide RNAs (sgRNAs) for the *cfap53* or *mns1* gene were designed using the web tool CHOPCHOP (https://chopchop.cbu.uib.no/). The target sites of sgRNAs were designed by identifying sequences that correspond to GGN(18)NGG in the DNA. sgRNAs were synthesized based on a modified two-component system ^46^. In brief, PCR was conducted to generate a DNA template containing the T7 polymerase binding site, the sgRNA target sequence, along with a common reverse oligonucleotide of sgRNA sequence. Phusion polymerase (NEB, M0530S) was employed for this process. Subsequently, sgRNA *in vitro* transcription was carried out using 500 ng of the purified DNA template, with the MEGAshortscript T7 kit (Ambion, AM1354M). 1 nl of a mixture of 700 ng of the Cas9 protein (Toolgen) and 700 ng of each sgRNA was injected into one-cell stage zebrafish embryos to generate F0 mutant fishes. The F0 fishes were outcrossed to wild-type and transmitting founders identified from embryo PCR and sequencing. Double mutants were generated using inter-crossing of the single mutant heterozygous fish.

### Mice

Mice were maintained at the Animal Facility of Riken Centre for Biosystems Dynamics Research (BDR), Japan, under a 12h light, 12h dark cycle and were provided with food and water *ad libitum*. All mouse experiments were approved by the Institutional Animal Care and Use Committee (permission number: A2016-01-6) and carried out in accordance with guidelines of the RIKEN BDR. *Mns1* knockout mice, lacking exon 3, was generated with the CRISPR-Cas9 system. Construction of the *Cfap53* mutant mouse strain has been described previously ^12^.

### Antibodies

Primary antibodies used in mouse study are rabbit anti-GFP (Invitrogen, A-11122), mouse anti-acetylated Tubulin (Sigma, T6793), mouse anti-γ-Tubulin (Sigma, T5326), rabbit anti-Mns1 (Sigma-Aldrich Prestige Antibodies, HPA039975). Antibodies used in the zebrafish study are rabbit anti-HA (Santa Cruz, SC805), rabbit anti-Myc (Santa Cruz, SC289), mouse anti-HA (Santa Cruz, SC7392), rabbit anti-GFP (Torrey Pines, TP401), mouse anti-acetylated-tubulin (Sigma, T6793), mouse anti-γ-tubulin (Sigma, T6557), anti-mouse HRP conjugate (Promega, W4028) and anti-rabbit HRP conjugate (Promega, W4018). Alexa Fluor-conjugated secondary antibodies for immunofluorescence staining were purchased from Invitrogen.

### Co-immunoprecipitation and western blot

Plasmid combinations of interest were transfected into HEK293T cells in 6 cm dishes, with each plasmid at a dosage of 3 µg per dish, utilizing Lipofectamine 2000 (Thermo Fisher Scientific). Following a 24-hour incubation period, the transfected cells underwent lysis in 800 µl of RIPA buffer (Thermo Fisher Scientific) supplemented with complete, Mini, EDTA-free Protease Inhibitor Cocktail (Roche, 11836170001). The resulting cell lysates underwent brief sonication and centrifugation. An aliquot from the clear cell lysate was extracted and subjected to boiling in 1× SDS loading buffer as the input (TCL). The remaining lysate was rotated overnight with 50 µl of Protein A-agarose beads (Roche) and 2 µg of mouse anti-HA (monoclonal, Santa Cruz, SC7392) or anti-myc (Santa Cruz, SC289) antibody. The beads were subjected to four washes in the immunoprecipitation (IP) buffer and then boiled in 50 µl of 1× SDS loading buffer (IP:HA). Both the TCL (30 µl) and IP (30 µl) fractions were separated using SDS-PAGE gels, transferred to PVDF membranes, blocked in 3% BSA, and probed with relevant primary antibodies (rabbit anti-HA (Santa Cruz, SC805) and rabbit anti-Myc (Santa Cruz, SC289)) and secondary antibodies (anti-mouse HRP conjugate (Promega, W4028) and anti-rabbit HRP conjugate (Promega, W4018)).

### Immunofluorescence staining of mouse embryos and trachea

Immunofluorescence staining of node cilia of E8.0 embryos and tracheal cilia was performed as described previously (Ide et al 2020, PlosGen). All antibody labelled samples were observed with an Olympus FV1000 and FV3000 microscope system.

### Whole-mount immuno-fluorescence staining of zebrafish embryos

Embryos were fixed for 2 hours at room temperature using Fish Fix (4% (w/v) paraformaldehyde and 4% (w/v) sucrose in PBS base). Subsequently, the fixed embryos were stored in methanol at -20°C. For immunofluorescence studies, embryos underwent a series of washes in decreasing methanol:PBS gradient, followed by PBS washes and blocking in PBDT (1% (w/v) BSA, 1% DMSO, 0.5% Triton X-100, PBS base) for 1 hour. Primary antibodies were introduced into PBDT and incubated with the embryos overnight at 4°C. Subsequently, embryos were washed in PBDT before incubating with fluorophore conjugated secondary antibodies and DAPI for 3 hours at room temperature. Stained embryos were cleared in 70% glycerol, mounted, and imaged using an Olympus Fluoview Upright Confocal Microscope. Image acquisition and analysis were performed using the Olympus Fluoview FV10-ASW software.

### Immunofluorescence analysis of HEK293T cells with MNS1 and CFAP53 overexpression

HEK293T cells were split and grown on glass coverslips in six-well plate one day before transfection. 2µg MNS1-HA/2µg MNS1-Myc and 2µg CFAP53-HA/2µg CFAP53-Myc plasmids were incubated with 20µl of Lipofectamine 2000 (Invitrogen) in 300 µl Optimal MEM (Invitrogen) for 5 min and added into cell culture medium. 18 hours after transfection, the transfected cells were fixed with 4% paraformaldehyde in PBS base for 30min. Rabbit anti-HA and mouse anti-Myc antibodies were used to stain the tagged proteins. Stained cells were mounted and imaged using an Olympus Fluoview Upright Confocal Microscope. Image acquisition and analysis were performed using the Olympus Fluoview FV10-ASW software.

### Live imaging with high-speed video microscopy

For high-speed video microscopy capturing cilia motility, zebrafish embryos were immobilized in 2% agarose (supplemented with 0.0175% Tricaine for 48 hpf embryos) and placed on 50 mm glass-bottom dishes. Ciliary motility was observed using a Zeiss 63X water-dipping objective mounted on an upright Zeiss AxioImager M2 microscope, which was equipped with an ORCA-Flash4.0 V2 C11440-22CU camera (Hamamatsu). Video processing and kymograph analysis were carried out using ImageJ 1.44d (NIH, USA). Mouse embryos were collected into DMEM-HEPES with 10% Fetal bovine serum (FBS). Motility of cilia was examined at 25°C with a high-speed CMOS camera (HAS-500M or HAS-U2, DITECT) at a frame rate of 100 frames/s for node cilia. The pattern of ciliary motion was traced and analysed with ImageJ and Photoshop CC (Adobe).

### TEM

E8.0 mouse embryos and adult mouse trachea were fixed with 2% paraformaldehyde (PFA) and 2.5% glutaraldehyde (GA) in 0.1 M phosphate buffer pH 7.4, 4°C. The samples were then washed with 0.1 M phosphate buffer and exposed to 2% osmium tetroxide. Ultrathin (70 nm) sections were prepared with a diamond knife, mounted on 200-mesh copper grids, stained with 2% uranyl acetate and lead stain solution, and observed with a JEM-1400 plus microscope (JEOL). Preparation of samples and imaging were performed in the facility of Tokai Electron Microscopy, Inc., Japan.

## Figure assembly

All figures were assembled using Adobe Illustrator CS4.

## Figure legends

**Supplementary Figure 1.**
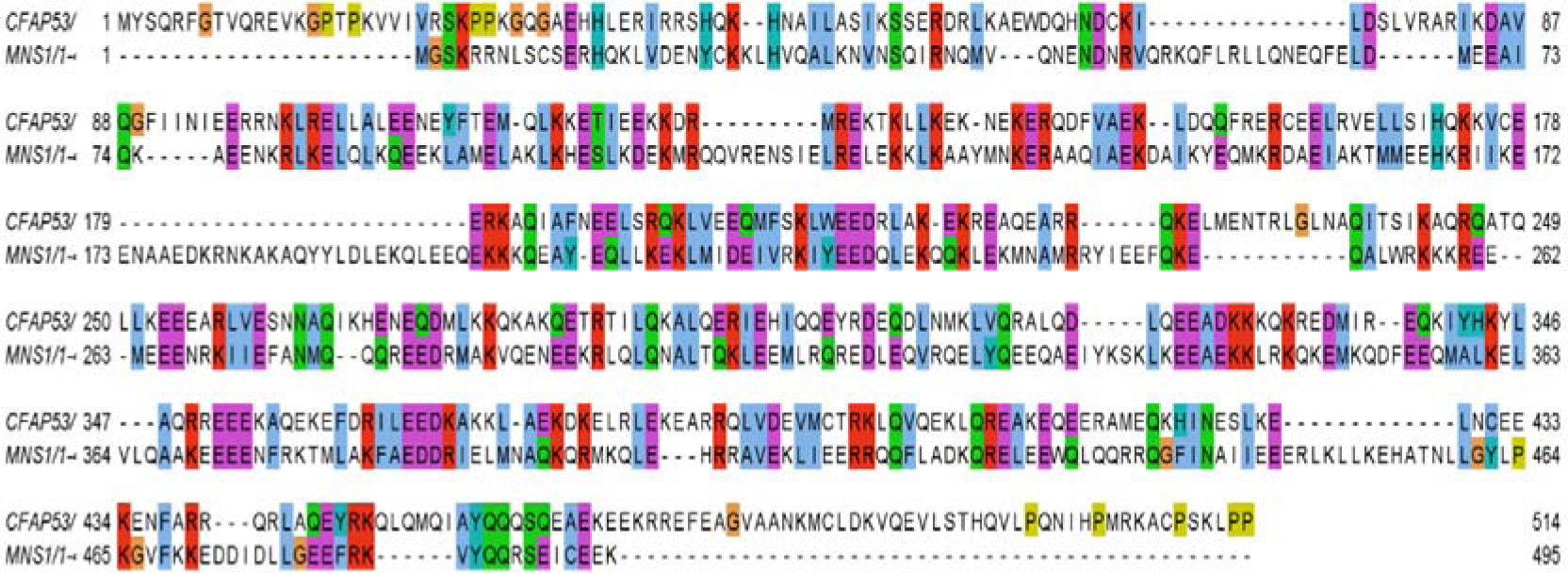
Global alignment of CFAP53 and MNS1 proteins from humans.

**Supplementary Figure 2.**
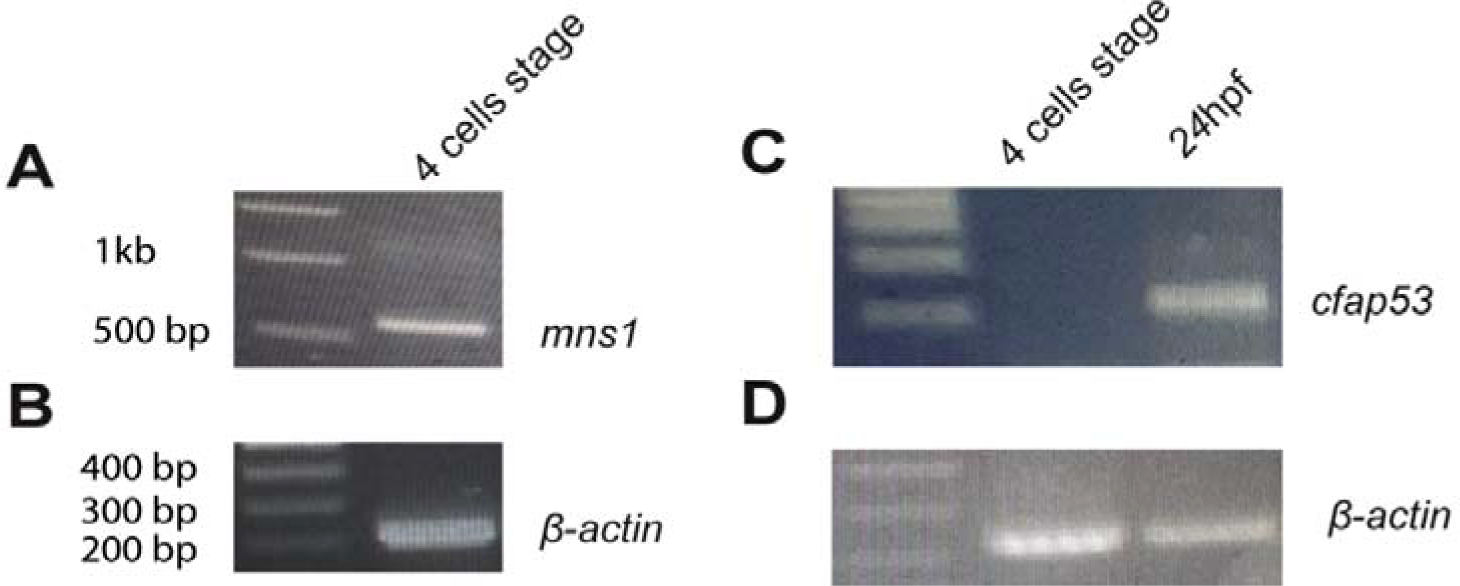
*mns1* mRNA is maternally deposited. Maternally deposited *mns1* mRNA was detectable in 4 cells stage zebrafish embryos by PCR, whereas *cfap53* mRNA was not detectable. Cfap53 mRNA could be detected by PCR using the same primer pair at 24 hpf. PCR of *βactin* was used as loading control.

**Supplementary Figure 3.**
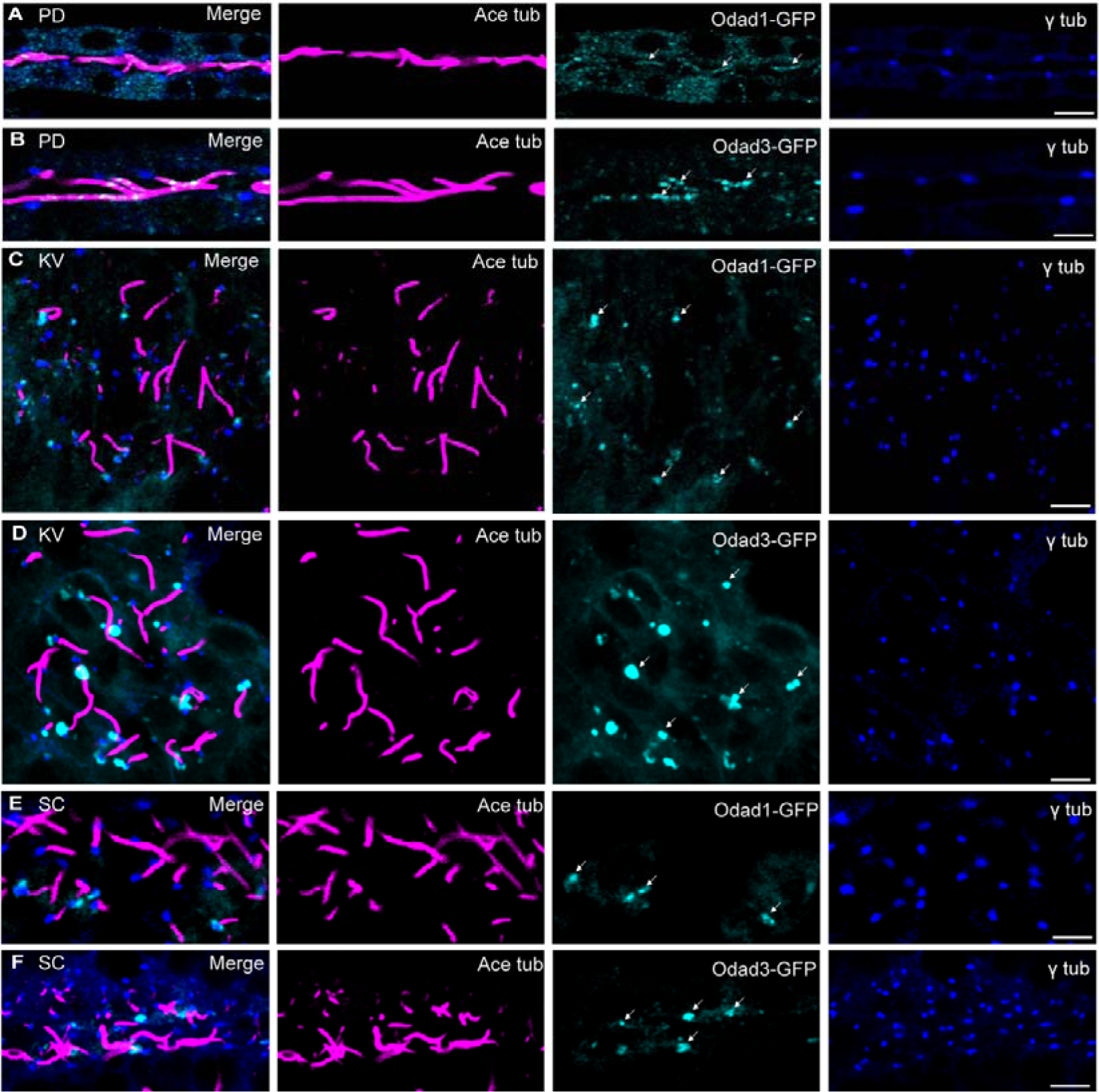
Localization of Odad1 and -3 proteins to motile cilia in zebrafish. (A, B) Odad1 and -3, tagged at the C-terminus with GFP, localized to the axoneme of (9+2) cilia in zebrafish pronephric duct (PD) at 24 hpf. (C, D) Odad1 and -3, tagged with GFP at the C-terminus, localized at the base of zebrafish KV cilia, close to and overlapping with the basal body (γ tubulin staining) at the 10 somites stage. (E, F) Odad1 and -3, tagged at the C-terminus with GFP, localized at the base of zebrafish spinal canal (SC) cilia, close to and overlapping with the basal body (γ tubulin staining) at 24 hpf. Scale bars = 5 µm. Localization studies were conducted in 3 technical replicates.

**Supplementary Table 1:** Sequence similarity between Mns1 and Cfap53 in selected species (pairwise alignments)

|  | *Rattus norvegicus* | *Mus musculus* | *Ornithorhynchus anatinus* | *Sarcophilus harrisii* | *Monodelphis domestica* | *Homo sapiens* |
| --- | --- | --- | --- | --- | --- | --- |
| **Gene Sequence Identity (%)** | 81.3 | 75.3 | 44.5 | 61.5 | 71.1 | 63.1 |
| **Protein Sequence Identity (%)** | 20 | 23.3 | 18.2 | 20 | 21.6 | 22.8 |

**Supplementary Table 2:**
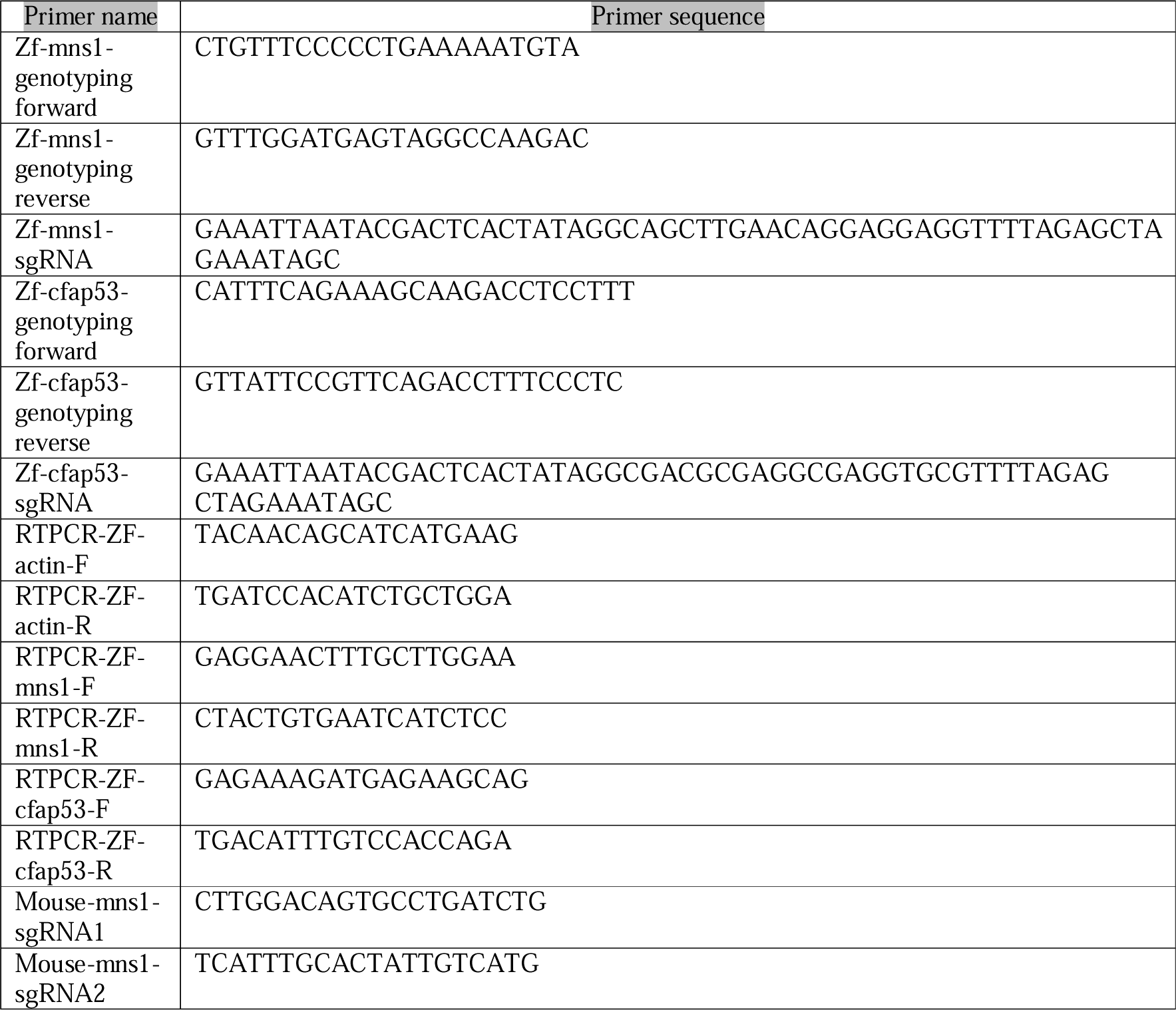
Primer and single guide RNA (sgRNA) sequences.

**Supplementary video 1. Motility of KV cilia of a wild-type zebrafish embryo.**

Rotary motion of the cilia is clearly visible. In this and all subsequent videos of KV cilia motility, embryos imaged were at the 10 somites stage (14 hpf). Videos were captured at 80 frames/sec (fps) and is played at 30 fps.

**Supplementary video 2. Motility of KV cilia of a mz *mns1* ^-/-^** zebrafish embryo. KV cilia were almost completely immotile.

**Supplementary video 3. Motility of nasal cilia of a wild-type zebrafish embryo.** Coordinated metachronal beating of multiple cilia on MCCs of the nose. In this and all subsequent videos of nasal cilia motility, embryos imaged were at 72 hpf. Videos were captured at 80 fps and is played at 30 fps.

**Supplementary video 4. Motility of nasal cilia of a mz m *ns1*^-/-^ zebrafish embryo**. Note a wild-type like pattern of ciliary motility.

**Supplementary video 5. Motility of node cilia of a *Mns1*** ^+/-^ **embryo.** The node cilia were motile. In this and all subsequent videos of the mouse node and tracheal cilia, the videos were captured at 80 fps and is played at 30 fps.

**Supplementary video 6. Motility of node cilia of a *Mns1* ^-/-^ embryo**. The node cilia were immotile.

**Supplementary Video 7. Motility of tracheal cilia of a *Mns1* ^+/+^ mouse.** The tracheal cilia were motile.

**Supplementary Video 8. Motility of tracheal cilia in a *Mns1*** ^-/-^ **mouse.** The tracheal cilia were motile, but they beat with smaller amplitude and frequency.

**Supplementary video 9. Motility of KV cilia of a *cfap53*^-/-^ zebrafish embryo.** KV cilia showed strong motility defect resembling *mns1* mutants.

**Supplementary video 10. Motility of nasal cilia of a *cfap53*^-/-^ zebrafish embryo**. Note a wild-type like pattern of ciliary motility.

**Supplementary video 11. Motility of nasal cilia of a zygotic *mns1*** ^-/-^**; *cfap53***^-/-^ **double mutant zebrafish embryo.** Nasal cilia exhibited aberrant motion, distinct from the smooth wave form characteristic of the wild-type embryo. Many cilia were completely paralyzed.

**Supplementary video 12. Severe motility defect of nasal cilia of a zygotic *mns1* ^-/-^; *cfap53* ^-/-^ double mutant embryo.** Majority of the nasal cilia were immotile. Remaining cilia exhibited very severe motility defect.

**Supplementary video 13. Motion of KV cilia in an *odad3* morpholino injected embryo.** KV cilia were immotile or highly dysmotile in the *odad3* morpholino injected embryo.

**Supplementary video 14. Motion of KV cilia in an *odad3* morphant embryo injected with *odad3* mRNA**. *odad3* mRNA significantly rescued KV cilia motility in *odad3* morpholino injected embryos.

